# A method for deep and quantitative protein profiling of urine sediment, soluble and exosome fractions for biomarker research

**DOI:** 10.1101/2023.06.26.546632

**Authors:** Peter Pichler, Amelie Kurnikowski, Manuel Matzinger, Karl Mechtler

## Abstract

Urine collection is painless and offers the potential for kidney liquid-biopsy(1), which appears particularly appealing with regard to the diagnosis of kidney disease (2) and patient follow-up after renal transplantation (3). From a nephrological point of view, urinary sediment and the soluble and exosome fractions of urine constitute different biological entities. We here describe a method that allows deep profiling of the protein content of the above-mentioned three fractions of urine by quantitative data-independent label-free proteomics. The method was evaluated using 19 urine samples from the Nephrology outpatient clinic at Vienna General Hospital, comprising a diverse set of chronic kidney disease (CKD) as well as patients after kidney transplantation (NTX). Peptide separation was accomplished through 60 min active gradients. A timsTOF Pro2 mass spectrometer was operated in DIA mode. The total analysis time per urine sample (three fractions) was around four hours. We demonstrate adequate technical and experimental reproducibility. Our data suggest that the protein information content of these three fractions is diverse, strengthening the importance of separate analysis. The depth of our quantitative proteomics approach permitted a detection of proteins characteristic for different parts of the nephron, such as Podocin, CD2-AP and Podocalyxin (Podocytes), SLC22A8 and SLC22A13 (proximal tubule) and Aquaporin-2 (collecting duct), suggesting that our method is sensitive enough to detect and quantify biologically relevant proteins that might serve as potential biomarkers. To the best of our knowledge, the ability to quantify up to 4000 protein groups per urine sample and more than 6000 protein groups in total makes our strategy the deepest proteome profiling study of urine to date. In conclusion, we established a method with promising figures of merit that we consider broadly applicable and useful for future clinical biomarker research studies in urine.

## Introduction

Patients attending the nephrology outpatient clinic typically provide blood and urine samples. Dozens of parameters are routinely measured in blood. Several of these parameters such as creatinine, blood urea nitrogen (BUN), serum electrolytes, and serum albumin and total protein provide prognostic and clinically relevant information to physicians. Notably, fewer prognostic parameters are routinely measured in urine. At the nephrology outpatient clinic of the Medical University of Vienna, urine is typically checked by dip-stick which measures glucose, protein, pH, nitrite, ketones, urobilinogen, blood, leukocytes and specific gravity. If dip-stick results are suggestive of urinary tract infection, urine culture is initiated. Upon physician request, urine samples are transferred to the central laboratory and the creatinine concentration as well as total protein and albumin concentrations are measured in urine. This permits calculation of the so-called urinary PCR and ACR (total protein to creatinine ratio and albumin to creatinine ratio). These are among the few biomarkers that are routinely measured in urine at present. Elevated PCR and/or ACR i.e. high amounts of protein in urine are indicate of potential kidney injury and high PCR and ACR in renal patients are associated with severe kidney disease and rapid progression to renal failure. (4, 5)

Urine collection is simple, rapid and painless. As kidney diseases may lead not only to general proteinuria and albuminuria but also to altered patterns in the urine proteome and peptidome, mass spectrometry-based peptidomics and proteomics have been used in the quest for kidney biomarkers in urine.(6) Capillary electrophoresis of urinary peptides coupled to quantification in MS1 scans has been applied successfully in this way. (2, 7–9) Bottom-up proteomics methods using LC-MS/MS offer the possibility of simultaneous identification and quantification of peptides and proteins, and such methods have also identified potential biomarkers (3, 10–12). However, so far few if any such biomarker peptides or proteins have made their way into routine workflows in the clinic.

Recent developments in mass spectrometry instrumentation and data processing have led to remarkable improvements in sensitivity and speed of data acquisition.(13, 14) We here evaluate the performance characteristics of quantitative data-independent label-free proteomics using a U3000 RSLC (Thermo Fisher Scientific) coupled to a timsTOF Pro2 mass spectrometer (Bruker Daltonics). To the best of our knowledge, this is the first urine proteomics study where urine is prefractionated into the three clinically relevant fractions urinary sediment, soluble urine and exosomes,(15) prior to further sample processing and measurement. In many previous urinary proteomics studies, urine is first centrifuged for 5–10 minutes at 400-1,000 g and only the supernatant is processed further. However, urine sediment is considered a rich source of information to the nephrologist. For this reason, we decided to analyze rather than discard urinary sediment. Recommendations for centrifugation such as the guidelines of the European urinalysis guidelines of the European Confederation of Laboratory Medicine recommend centrifugation at 400 g for 5 minutes (16), which is sufficient for visual inspection of urinary sediment. However, higher g and longer centrifugation time is required to sediment all particulate components of urine.(17) To combine precipitation of cellular urine components and urinary casts with a removal of bacteria and cell debris, we decided on centrifugation at 2,000 g for 30 min. Of note, this also corresponds to the recommendations in the Total Exosome Isolation kit.

## Methods

### Sample collection

Urine samples were collected from 19 patients upon appointment at the nephrology out-patient clinic of Vienna Medical University, after informed consent had been obtained. We intentionally obtained fresh spot urine (rather than 24 hour urine) to obtain samples similar to those provided by kidney patients on a general basis. Urine was stored at 4°C within a few minutes (maximum 30 minutes). Further storage was at -20°C within 24 hours and at -70°C within 2 weeks.(18)

Average age of patients was about 55 years, with a broad range from 27 to 77 years. More than half of the patients had undergone kidney transplantation (NTX). As in many nephrology studies, male patients were somewhat over-represented. The small cohort comprised patients with a broad range of nephrologic diseases and suspected causes for CKD (chronic kidney disease) such as diabetes mellitus and arterial hypertension, autosomal dominant polycystic kidney disease (ADPKD), various types of glomerulonephritis, monoclonal gammopathy, aHUS (atypical Hemolytic Uremic Syndrome) and rare causes. Around a fourth of patients received SLGT2i medication. The diverse cohort was chosen to evaluate and establish the broad applicability of our method to urine analysis in patients with various types of kidney disease as well as post kidney transplantation.

### Preparation of three urine fractions sediment, soluble urine and exosomes

Urine was thawed in a water bath (37°) and centrifuged at 2,000 g for 30 min at room temperature. Supernatant was collected: this constituted the “soluble urine fractions”, which were precipitated over-night in 4x volume of cold acetone at -20°C. The remaining sediment pellets were washed in phosphate-buffered saline and spun for 20 min at 2,000 g at room temperature. Washing was performed twice. Pellets were resuspended in 50–100μL of 8 M Urea 100 mM Tris pH 7.5, vortexed for 10s and sonicated for 1 min. Disulfide bridges were reduced in 5 mM DTT (dithiothreiotol) for 30 min at 37°C and thiol groups were alkylated in 20 mM IAA (iodoacetamide) for 30 min in the dark. Samples were quenched by adding 2.5 mM DTT and diluted to 6 M Urea in 100 mM Tris pH 7.5. Benzonase 125 IU was added, followed by protease LysC at an estimated amount of 1:50 (weight/weight) and digestion was performed for 2 hours at 37°C. Subsequently, Trypsin 1:50 w/w was added and digestion of “sediment fraction” samples was continued at 37°C over-night. Digestion was stopped by adding 10% TFA to a final concentration of 1%. On the following day, acetone precipitated “soluble fraction” samples were centrifuged at 4°C for 30 min at 20,000 g. Protein pellets were resuspended in 50-100μL of 8M Urea 100mM Tris pH 7.5, vortexed and sonicated for 2 min. Further sample processing was akin to “sediment” fractions, except no benzonase was used.

To prepare exosomes, 2.5 mL of “soluble urine” fraction (supernatant obtained after 30 min centrifugation of urine at 2,000 g) was transferred to 5 mL low binding Eppendorf tubes, and 2.5 mL of “Total Exosome Isolation” reagent (Thermo Fisher Scientific) was added. After vortexing and incubation for 1 hr at room temperature, samples were centrifuged at 4°C for 1 hr at 20,000 g. Pellets containing exosome vesicles might not be visible at this point. The supernatant was removed completely. Vesicles were resuspended in 50 uL of 100 mM glycine pH 7.55 8 M Urea 5% SDS, vortexed for 10s and sonicated for 1 min. 25 μL of each sample was reduced in 5 mM DTT, alkylated in 20 mM IAA and quenched by adding 2.5 mM DTT. Samples were acidified by adding 2.5 μL of 55% phosphoric acid. 165 μL of S-trap binding buffer (90% MeOH, 100 mM TEAB pH 7.55) was added to each S-trap micro column. Then 2 μg of Trypsin/LysC solution (1 μg/μL each) was added to each acidified sample, immediately mixed by pipetting up and down, transferred to the S-trap, and mixed again by pipetting up and down. S-trap columns were placed into 1.7 mL Eppendorf tubes and centrifuged for 30s at 10,000 g. S-trap columns were washed three times in 150 μL S-trap buffer, discarding the flow-through, as described in the S-trap protocol. Trypsin 1 μg in 25 μL of 50 mM TEAB pH 8 was added to each S-trap column. Caps were closed loosely, and samples were incubated for 2hr at 47° in water-saturated atmosphere. After digestion, peptides were eluted first with 40 uL of 50 mM TEAB, second 0.2% formic acid and third 50% acetonitrile 0.2% formic acid, each time followed by centrifugation for 30s at 10,000 g. Flow-through was collected, 10% TFA was added to a final concentration of 1%. Samples were concentrated (without drying completely) on a SpeedVac.

### Data acquisition and processing

200 ng of each fraction (sediment, soluble urine and exosomes) was injected on a U-3000 RSLC Nano-HPLC system (Thermo Fisher Scientific) coupled to a timsTOF Pro2 or timsTOF HT mass spectrometer (Bruker Daltonics). Precolumn loading was at a flow rate of 10 μL/min, with 0.1% TFA mobile phase. An IonOpticks Aurora 25 cm column (generation 2) was used for peptide separation. Mobile phases were A: 0.1% FA and B: 0.08% FA in 80% acetonitrile. The precolumn was switched into the column flow at 4.0 min. The LC gradient was as follows: 4 min: 2% B, 49 min: 24% B, 64 min: 35% B, then 95% B for 5 min and equilibration with 2% B until 80.0 min.

The timsTOF Pro2 was operated in DIA mode: 1 MS1 scan was followed by 8 DIA-PASEF frames: frame1: 1/k0 1.01-1.42 m/z 800-825, 1/k0 0.83-1.01 m/z 600-625, 1/k0 0.64-0.83 m/z 400-425; frame2: 1/k0 1.04-1.42 m/z 825-850, 1/k0 0.85-1.04 m/z 625-650, 1/k0 0.64-0.85 m/z 425-450; frame3: 1/k0 1.06-1.42 m/z 850-875, 1/k0 0.87-1.06 m/z 650-675, 1/k0 0.64-0.87 m/z 450-475; frame4: 1/k0 1.09-1.42 m/z 875-900, 1/k0 0.90-1.09 m/z 675-700, 1/k0 0.64-0.90 m/z 475-500; frame5: 1/k0 1.11-1.42 m/z 900-925, 1/k0 0.92-1.11 m/z 700-725, 1/k0 0.64-0.92 m/z 500-525; frame6: 1/k0 1.13-1.42 m/z 925-950, 1/k0 0.94-1.13 m/z 725-750, 1/k0 0.64-0.94 m/z 525-550; frame7: 1/k0 1.16-1.42 m/z 950-975, 1/k0 0.97-1.16 m/z 750-775, 1/k0 0.64-0.97 m/z 550-575; frame8: 1/k0 1.18-1.42 m/z 975-1000, 1/k0 0.99-1.18 m/z 775-800, 1/k0 0.64-0.99 m/z 575-600). Accumulation and ramp time were set to 166 ms. Collision energy was 20 eV at 1/k0 0.6 Vs/cm^2^, and 80 eV at 1/k0 1.6 Vs/cm^2^ Automatic calibration was set on.

DIA-NN 1.8.2 was employed for data processing. Library-free searches were performed using swissprot human reference proteome fasta downloaded on 27 May 2022, supplemented with “Universal Protein Contaminants” fasta.(19) One missed cleavage was permitted, and N-terminal excision of Methionine and carbamidomethylation of Cysteine were set as modifications. The final database comprised 20800 proteins from which 4333185 precursors were generated by DIA-NN. Match between runs was set off, precursor and protein group false discovery rate were 1%, protein inference was set to isoform IDs, and heuristic protein inference was enabled to ensure maximum parsimony results (for benchmarking purpose).

## Results

### Time requirement of workflow

The sample preparation workflow is depicted in Fig.1. It is feasible to complete the workflow within 1.5 days. After diluting sediment and soluble urine samples to 6 M urea, in-solution digestion with LysC is performed for two hours. Subsequently, urea concentration is brought to 2 M and Trypsin is added for over-night digestion. Exosome sample preparation can even be performed within a single workday, as the time required for digestion on S-trap columns is only 2 hours.

**Fig. 1.**
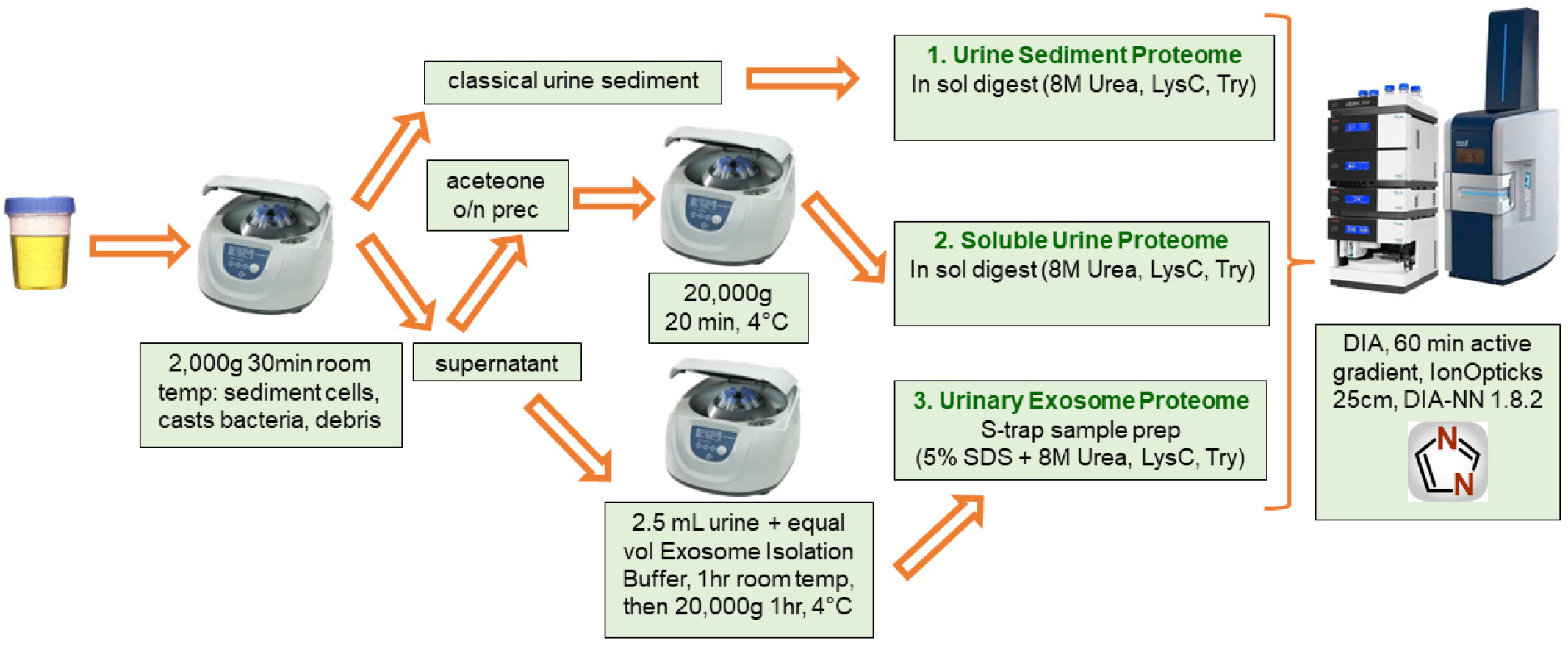
Experimental Workflow. Urine was collected in the outpatient clinic and stored at 4°C within 30 minutes. Urine was centrifuged at 2,000g for 30 min. Centrifugation speed and time exceeded settings typically used to generate urinary sediment for visual inspection, as our goal was to generate supernatant free of cells, casts, bacteria and debris. Urine sediment was processed by solubilization in 8 M urea and sonication, followed by in solution digestion with LysC and Trypsin. Part of the supernatant was acetone precipitated over-night, with further sample processing akin to sediment. In addition, 2.5 mL of supernatant and 2.5 mL of Total Exosome Isolation reagent were mixed and incubated at room temperature for 1 hr, followed by centrifugation at 4°C for 1hr at 20,000 g. The pellet was resolubilized in S-trap buffer containing 5% SDS and 8 M Urea. Further sample processing was according to the S-trap “high recovery” sample preparation protocol, which ensures digestion of proteins captured by the S-trap as well as removal of PEG and SDS. Typically, 200 ng of each sample was injected on a U-3000 RSLC. Active gradient time was 60 minutes, and an IonOpticks Aurora 25cm column was used for peptide separation. Three samples (sediment, soluble fraction, exosomes) were measured in DIA mode on a timsTOF Pro2 instrument. DIA-NN 1.8.2 was used for data processing.

Total analysis time for the three fractions of a urine sample was approximately 4 hours (60 min active gradient, 85 min total LC time per sample), which should be compatible with a deployment of the method in clinical studies. The proportional Venn diagram in Fig.2 depicts the overlap between the numbers of identified and quantified protein groups in the three fractions sediment, soluble urine and exosomes from patient 2. Although a considerable part of the protein groups is detected in more than one fraction, the scatter plots in Fig.2 illustrate a low to only moderate correlation of quantitative abundance between fractions. This suggests that separation of urine into three fractions sediment, soluble urine and exosomes is indeed meaningful and that separate measurement of these three fractions increases the information gained through the experiment.

**Fig. 2.**
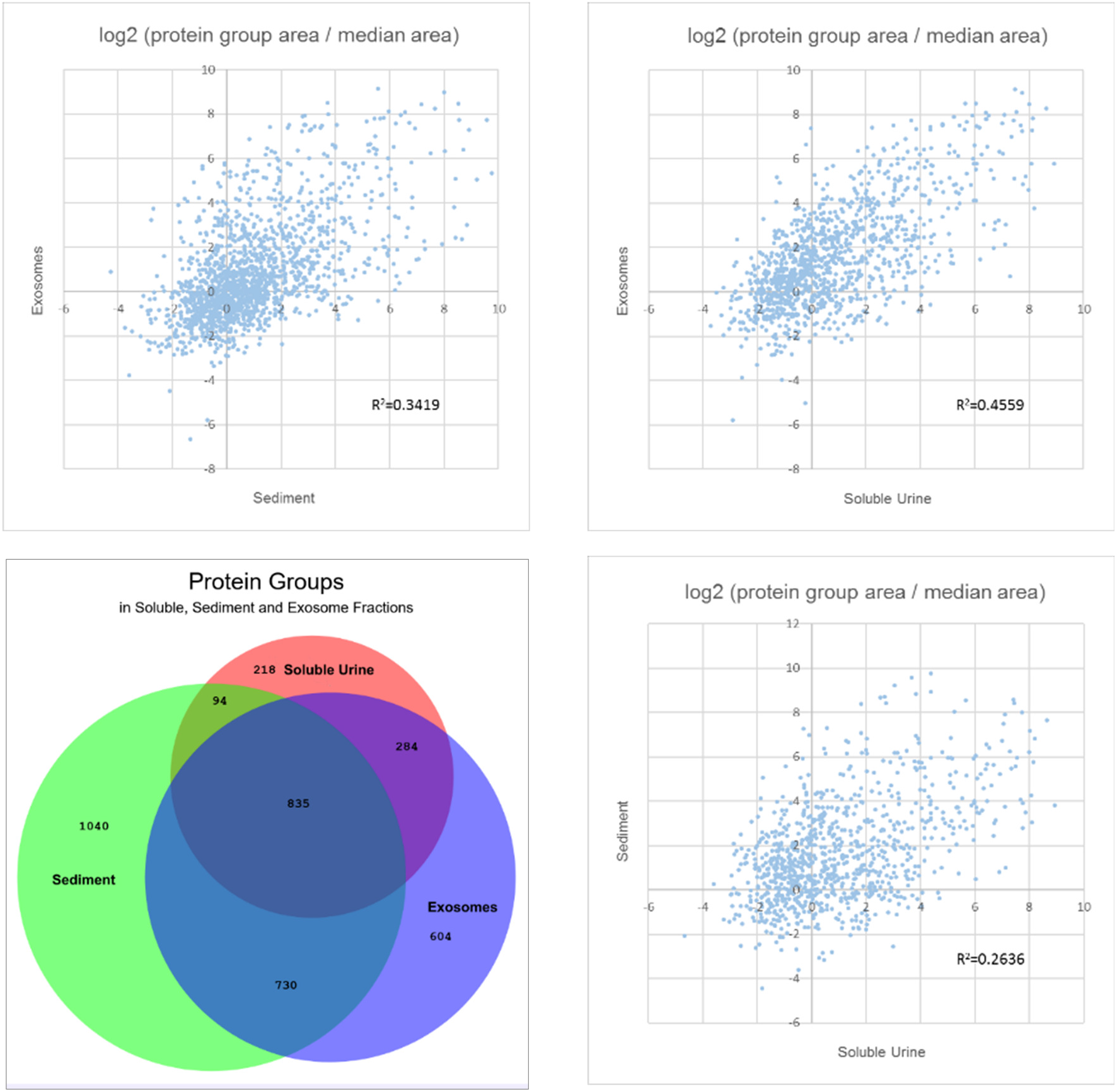
Proportional Venn Diagram. (30) illustrating the number of proteins groups detected and quantified in the three fractions sediment, soluble urine and exosomes from patient 2 (lower left). In total, 3805 protein groups were detected and quantified in this urine sample. **Scatter Plots** depicting the correlation of quantitative abundance between the protein groups detected in common in fractions sediment+soluble, soluble+exosomes and sediment+exosomes, respectively. Quantitative abundance was calculated for each protein group as the log2 transformed ratio of the specific protein group area to the median protein group area in the respective fraction.

### Initial quality control and quantification of protein amount after digestion

We routinely performed a first quality control check by injecting a small amount of each sample on a stand-alone LC-MS system equipped with a monolithic column and a UV-detector recording at 214 nm. A comparison of the area under the elution curve (AUC) of a 100 ng peptide standard from a commercially available HeLa digest (Thermo Fisher Scientific) to the AUC of each individual sample was used to estimate the amount of sample obtained after digestion (S-trap or in-solution). In addition, the recorded UV signal was inspected for potential time shifts of the elution profile which would suggest inadequate digestion, so that such samples, if any, could undergo a second round of digestion before measurement.

We also routinely evaluated the performance of the U-3000 timsTOF LC-MS system by monitoring figures of merit obtained with 100 ng HeLa digest standard, before analysis of any urine samples. In addition, blank samples were analyzed before and between urine samples, to exclude blending by carry-over from previous samples.

### Exosome sample preparation by Total Exosome Isolation reagent followed by S-trap protocol

Exosome sample preparation using Thermo Fisher Scientific “Total Exosome Isolation” reagent (previously Invitrogen / Life Technologies) had been previously shown to entail a risk of generating samples contaminated with polymer(20). From information in an exosome sample preparation patent held by Life Technologies(21), we presume that the kit might contain PEG-6000 or other types of PEG to encourage precipitation of exosome vesicles at 20,000 g. This is why we chose the S-trap protocol for protein digestion after exosome preparation, as the S-trap should remove both PEG and SDS. In addition, the S-trap protocol typically employs 5% SDS as a sample resuspension buffer, which appears beneficial for solubilization of proteins from vesicles.

To evaluate the effect of possible remnants of PEG or SDS on peptide separation, we first injected 100 ng of a commercially available HeLa peptide digest standard onto a stand-alone LC equipped with a monolithic column. Adding increasing amounts of PEG and SDS to this sample caused a time shift of peptide elution that we did not observe with our digested exosome samples (data not shown). In addition, measurement of 200 ng of exosome samples on our LC-MS system consisting of U-3000 and timsTOF did not unveil polymer clusters as described in (20). We thus consider our exosome preparation protocol (Total Exosome Isolation reagent followed by S-trap) amenable to LC-MS analysis.

### Technical and experimental reproducibility

Next, we evaluated the precision of data acquisition. Four injections of samples 11s and 17s respectively, were measured on a U-3000 RSLC coupled to a timsTOF Pro2 mass spectrometer (Fig.3). Results indicated adequate reproducibility with median CV approaching 10% for technical replicates from soluble urine samples. Of note, the numbers of protein groups identified from urine sediment were typically higher than from soluble urine, which might explain the somewhat higher median CV of up to 15% obtained with urine sediment samples 11p and 17p.

**Fig. 3.**
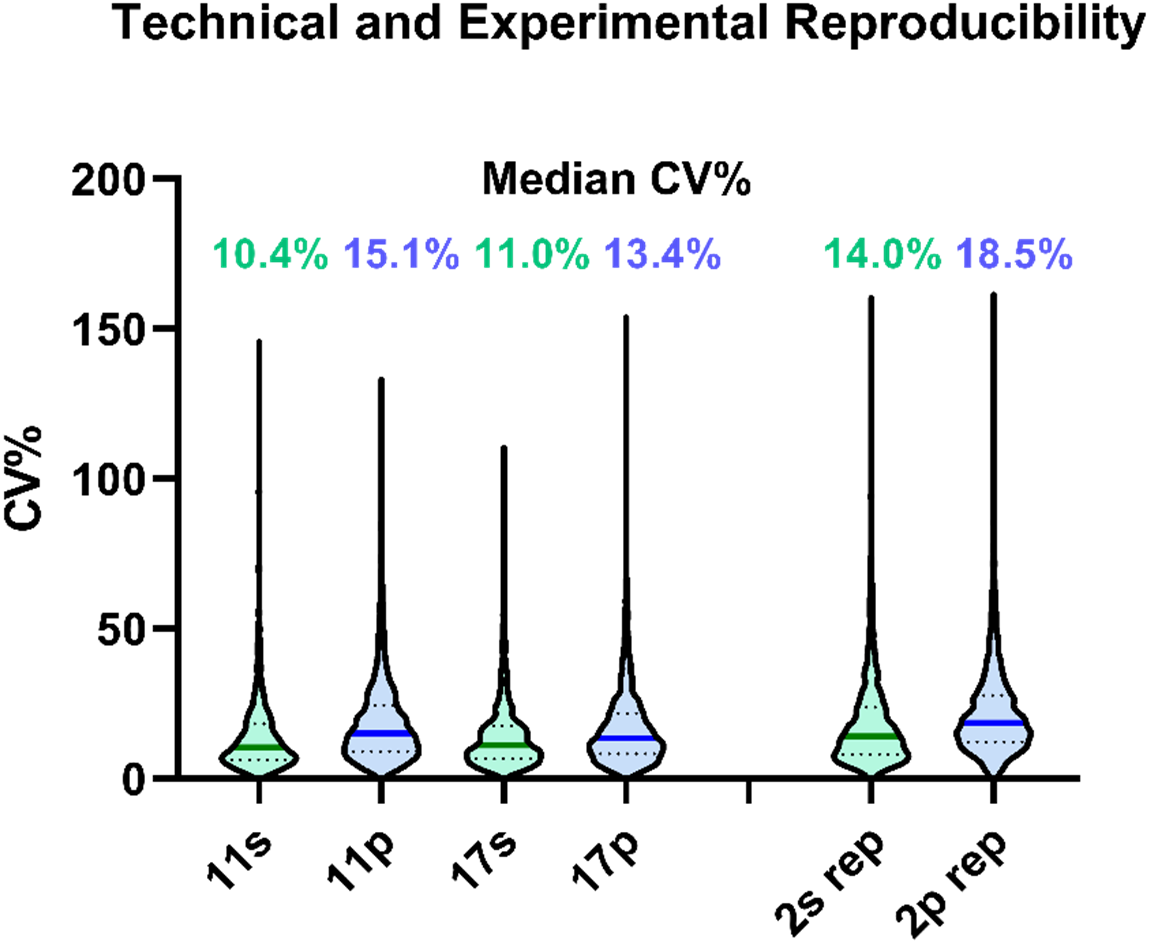
Technical and Experimental Reproducibility. Samples 11s+11p and 17s+17p (soluble and pellet/sediment) were measured four times to evaluate technical reproducibility: Violin plots illustrating the distribution of CV values for quantified protein groups are shown (left part). For comparison, the experimental sample workflow was performed four times to generate four samples 2s-rep and 2p-rep each, for an evaluation of experimental reproducibility (right part). Median CV is given above each violin plot.

For comparison, four experimental replicates of soluble urine and sediment (sample preparation workflow performed four times for each sample) showed experimental CV well below 20% for samples 2s-rep and 2p-rep (Fig.3).

As Nagaraj and Mann reported a median CV of 48% for intra-individual day-to-day variability of the urine core proteome, and a median CV of 66% for inter-individual variability between healthy individuals,(22) we consider reproducibility and precision of our method sufficient for biomarker discovery in urine.

### Data analysis: identified and quantified protein groups

Settings for data processing with DIA-NN 1.8.2 were as follows: no match-between runs, precursor and protein group false discovery rate 1%, protein inference: isoform IDs, heuristic protein inference enabled to ensure maximum parsimony. Box-and-whisker plots of the numbers of protein groups identified and quantified in the 19 urine samples with at least one unique peptide are depicted in Fig.4: The highest numbers of protein groups were identified in urinary sediment (median 2568 protein groups, range 992 to 4044). Exosomes provided the second highest numbers (median 2369 protein groups, range 1090 to 2453), and soluble proteins detected in supernatant after centrifugation the third (median 1382 protein groups, range 545 to 2053). Exosome sample preparation had been performed for samples 2, 11, 12 and 16. In total, between 1960 and 4390 protein groups (range) could be identified and quantified in the urine samples, with a median of 3080 protein groups per urine sample. In total, 6247 protein groups (and 66454 precursors) were identified in a combined DIA-NN search of all samples. To the best of our knowledge, this makes our study the deepest urine proteomics study to date.

**Fig. 4.**
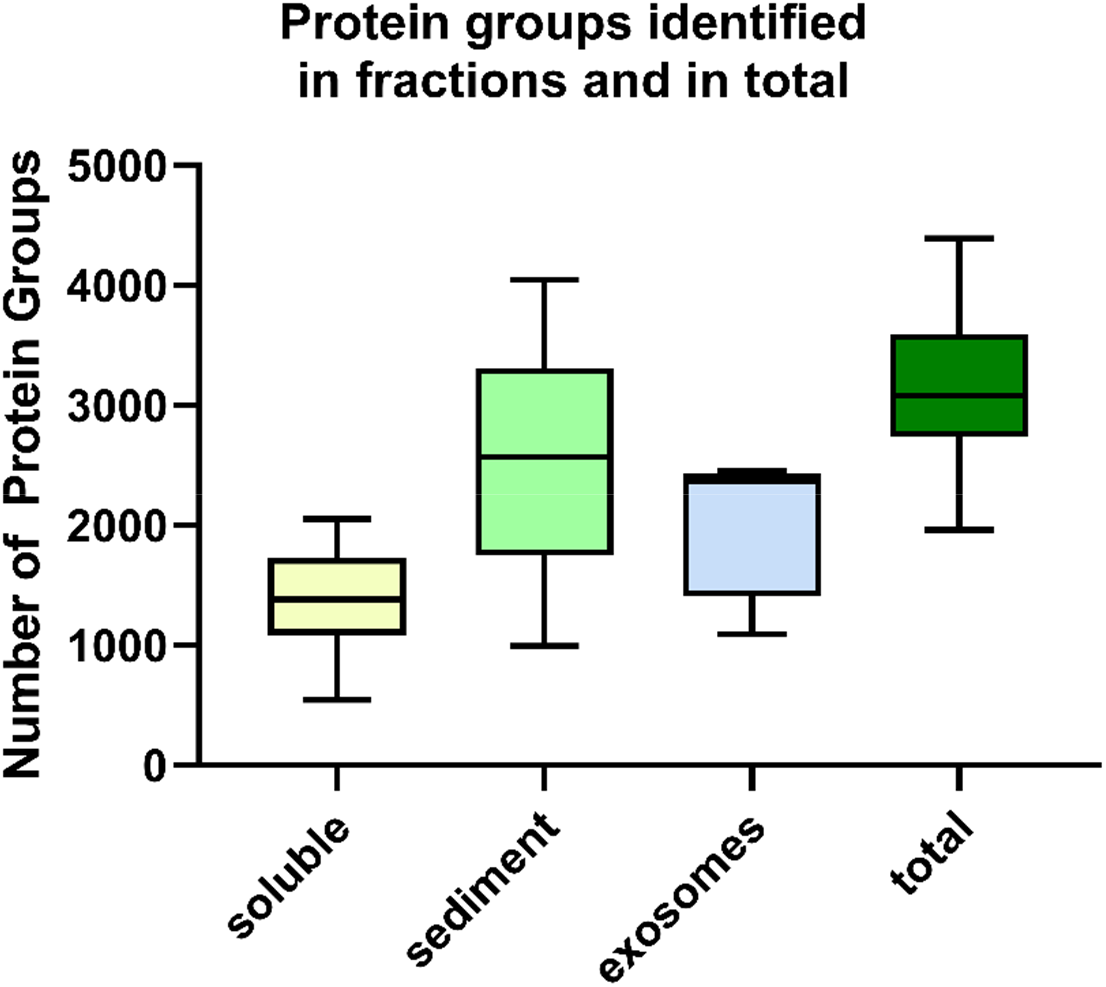
Number of Protein Groups. identified in the three fractions (soluble urine, sediment, exosomes) and in total (Box Plot, whiskers Tukey).

### Potential biological relevance

We further assessed whether the achieved depth of proteome coverage might permit identification and quantification of biologically relevant proteins:

Inspection of results indicate that our method is capable of detecting proteins characteristic for different parts of the nephron that play crucial biological functions in the respective cells they are derived from, such as Podocin, CD2-AP and Podocalyxin from Podocytes, SLC22A8 and SLC22A13 from proximal tubule cells, and Aquaporin-2 from collecting duct cells.

Podocytes are cells that line the visceral layer of Bowman’s capsule in the kidney glomerulus. Interdigitations of podocyte foot processes form the so-called “slit diaphragm”, a structure involved in the filtering of primary urine that is required to prevent proteinuria.(23) Podocytes express Podocin and Podocalyxin.(24) CD2-AP (CD2-associated protein) seems to be involved in anchoring the slit diaphragm to the podocyte cytoskeleton. It is known that loss of podocytes occurs early in patients developing diabetic kidney disease.(25) For this reason, we hypothesize that a detection of podocyte proteins as possible markers of podocyte injury might be of avail for the early diagnosis of diabetic kidney disease. The quantification of podocyte protein markers in urine might also help assess the prognosis and response to drugs that preserve kidney function in diabetic patients.

Of note, average total Podocin in the four samples from diabetic patients was by a factor of 2.46 higher than average total Podocin from three patients with ADPKD (Fig.5A), consistent with the notion that podocyte injury might play a more prominent role in diabetes than in ADPKD. It is known that treatment with SGLT2i medication preserves kidney function, in CKD patients with and without diabetes.(26) We therefore find it interesting that CD2-AP was 36% lower and Podocalyxin was 48% lower in diabetes patients receiving SGLT2i drugs compared to patients not receiving such drugs (Fig.5A). We are aware that these preliminary data are based on very low numbers of patients and might thus constitute chance observations. Still, the findings suggest that quantification of urinary Podocin, CD2-AP and Podocalyxin (and possibly other proteins associated with podocytes) might provide clues to early diagnosis of diabetic kidney injury or the treatment response to kidney-protecting drugs in future studies with larger numbers of patients.

**Fig. 5.**
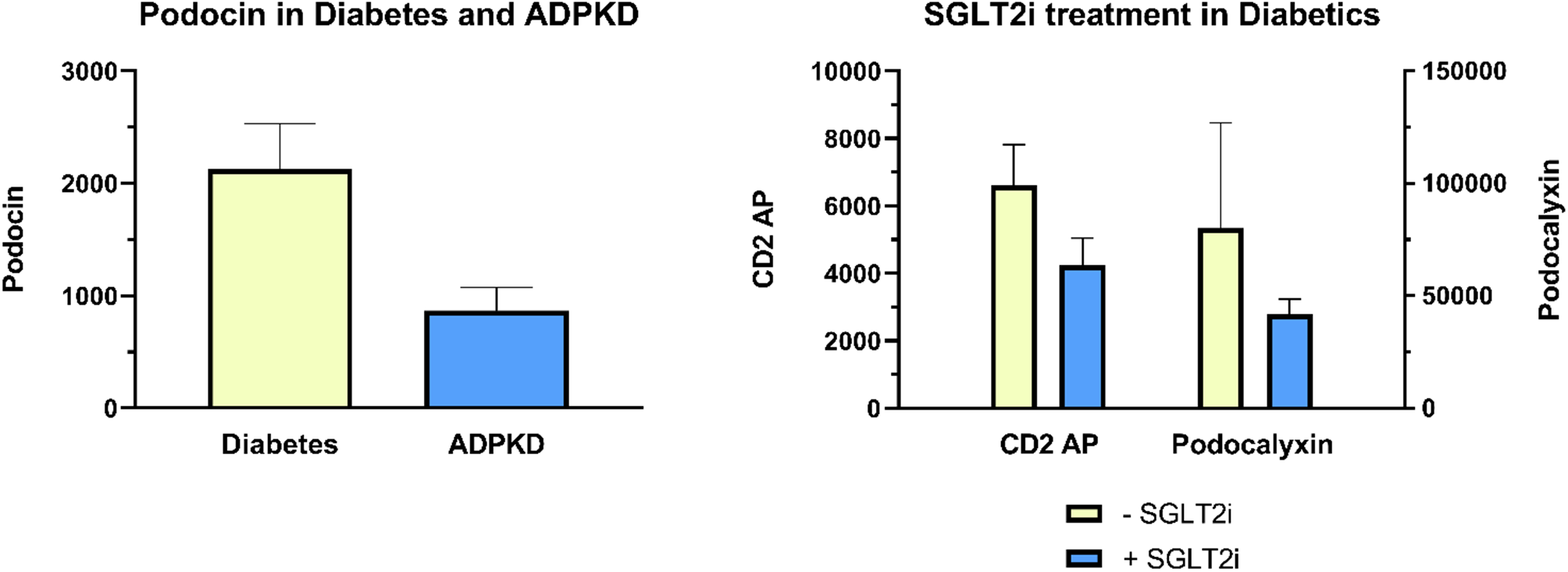
A. Quantification of Podocin in urine of diabetic (N=4) and ADPKD (N=3) patients. Of note, Podocin was detected in all diabetes patients but in only two out of three ADPKD patients. Even in the two ADPKD patients where Podocin could be detected, Podocin values were lower than in diabetes patients. B. **Response of CD2-AP and Podocalyxin** to treatment with SGLT2i in diabetic patients (N=4, half of whom received treatment).

In addition, the only soluble urine fraction where we were able to quantify PKD1 (Polycystin, the protein that causes polycystic kidney disease when mutated) was patient 1, a patient with known polycystic kidney disease patient who had not yet received a kidney transplant.

These hints to potential biological relevance of the proteins quantified by our method, together with the data on reproducibility and the ability to perform sample preparation and measurement within a moderate amount of time, suggest that our method may prove useful in future larger-scale clinical studies.

## Discussion

For application in a clinical setting such as a clinical study, sample processing and analysis should be simple and rapid.

Until a few years ago, deep sampling of the urinary proteome required extensive sample prefractionation e.g. by two-dimensional chromatography, ultracentrifugation, combinatorial peptide ligand libraries, lectin affinity or excision of bands from gel electrophoresis.(27–29) While these methods permitted a considerable depth of urine proteome coverage up to ca 3400 protein groups, such methods are difficult to implement in clinical studies, when large sample numbers are required to reach adequate statistical power. For comparison, urine sample preparation with our straightforward “three fractions” method can be completed in 1.5 workdays, with the possibility to process many urine samples in parallel. Measurement time is ca 4 hours including column equilibration (3 × 60 min active gradient time). And data analysis with DIA-NN can be accomplished in about the same time per sample, depending on the available computational power. The attained sensitivity and depth of analysis (median 3080 protein groups per urine sample, range 1960 to 4390) suggests that our method captures a considerable part of the information content of urine. Our strategy of dividing urine samples into three clinically relevant fractions i.e. urinary sediment, soluble urine and exosomes, should help preserve this information. We observed technical CVs of 10-15% (four repeated injections of the same sample) and experimental CVs < 20% (measurement of four samples generated by repeating the sample preparation workflow four times), indicating that the method is precise enough for urine analysis. For comparison, Nagaraj and Mann reported a median intra-individual day-to-day CV of 48% for the urine core proteome and a median inter-individual CV of 66%.(22)

In the quest for high numbers of identified and quantified protein groups, we do not consider mere numbers *per se* of importance, but rather whether the attained sensitivity permits identification and quantification of biologically relevant proteins, thus offering the possibility to answer biological questions. Our method proved capable of identifying proteins characteristic for different cells along the nephron, such as Podocin, CD2-AP and Podocalyxin (Podocytes), SLC22A8 and SLC22A13 (proximal tubule cells) and Aquaporin-2 (collecting duct). In addition, our data showed Podocin 2.46-fold elevated in diabetic as compared to ADPKD patients, and SGLT2i treatment seemed to reduce CD2-AP and Podocalyxin in patients with diabetes consistent with the kidney-preserving effect of these drugs. We hypothesize that quantification of the amount of these proteins in future large-scale clinical studies might lead to the detection of protein markers that aid clinicians in early detection and evaluation of the prognosis and drug response of kidney diseases, including diabetic kidney disease and the follow-up of patients after kidney transplantation.

We are aware of the limitations of our study in terms of the small numbers of patients and their heterogeneity. However, goals of this study were first to establish methodology for urine analysis with adequate figures of merit for future larger-scale clinical studies, and second to provide proof of concept that the established methodology appears applicable to a broad range of kidney diseases, including renal patients post NTX (kidney transplantation). Third, preliminary findings with regard to the above-mentioned biologically relevant proteins suggest that our method is capable of sampling the information content of urine to a depth that may allow detection of clinically relevant biomarkers in large-scale clinical studies.

Future variations of our method may employ shorter LC columns and shorter net gradient time and an automatization of sample preparation.

## Abbreviations used

CKD: chronic kidney disease
NTX: kidney transplantation
timsTOF: trapped ion mobility spectrometry Time-Of-Flight
DIA: Data Independent Acquisition
BUN: Blood Urea Nitrogen
PCR: urinary Protein Creatinine Ratio
ACR: urinary Albumin Creatinine Ratio
LC-MS: Liquid Chromatography Mass Spectrometry
MS1: survey scan for the detection of intact analyte m/z
MS/MS: m/z scan following analyte fragmentation in the gas phase to generate fragment spectra
RSLC: rapid separation LC system
aHUS: atypical Hemolytic Uremic Syndrome
SGLT2i: drugs that inhibit sodium-glucose cotransporter-2 in the proximal tubule of the nephron thereby increasing glucosuria and reducing blood glucose levels
DTT: Dithiothreitol
IAA: Iodoacetamide, Tris Trishydroxymethylaminomethan
TFA: Trifluoroacetic acid
SDS: Sodium dodecyl sulfate
MeOH: Methanol
TEAB: Triethylammonium bicarbonate
PEG: Polyethylene Glycol
FA: Formic Acid
ACN: Acetonitrile
PASEF: Parallel Accumulation-Serial Fragmentation
CV: Coefficient of Variation
DM: Diabetes mellitus
DM II: Diabetes Type 2
DM I: Diabetes Type I
aHT: arterial HyperTension
ADPKD: Autosomal Dominant Polycystic Kidney Disease
IgA GN: IgA Glomerulonephritis
aHUS: atypical Hemolytic Uremic Syndrome
TMA: Thrombotic MicroAngiopathy
CFI: Complement Factor I
CD2-AP: CD2-associated protein

## Acknowledgements

This work was supported by grant LS20-079 of the Vienna Science and Technology Fund https://hd-research.net/ All LC-MS/MS analyses were performed on instruments of the Vienna BioCenter Core Facilities instrument pool. The authors thank the IMP for general funding and access to infrastructure and Prof. Manfred Hecking (Medical University of Vienna, Department of Medicine III, Vienna, Austria) for funding of this specific project. We also thank Goran Mitulovic and Gary Kruppa (Bruker Daltonics, Bremen, Germany) for advice and support, and Karel Stejskal and Gabriela Krššáková (Proteomics Core Facility, IMP, Vienna, Austria) for continuous support and maintenance of instruments.

## Figures & Tables

Tables 1 and 2. **Patient characteristics and suspected causes of CKD**

**Table. 1.** Characteristics of the Patients.

| TABLE.1. **Characteristics of the Patients** |  |
| --- | --- |
| Age – yr (range) | 54.9 +/- 14.9 (range 27 - 77) |
| Female (%) | 7 (37%) |
| NTX (%) | 11 (58%) |
| DM I (%) | 1 (5%) |
| DM II (%) | 3 (16%) |
| SGLT2i treatment | 5 (26%) |

NTX: St.p. kidney transplantation, DM I: Diabetes mellitus Type I, DM II: Diabetes mellitus Type II, SGLT2i: treatment with SGLT2 inhibitor medication

**Table. 2.** Suspected cause of CKD in the Patients.

| TABLE.2. **Suspected cause of CKD in the Patients.** |  |
| --- | --- |
| DM +/- aHT | 4 |
| aHT only | 2 |
| ADPKD | 3 |
| IgA GN | 2 |
| monoclonal gammopathy | 2 |
| membranous GN | 1 |
| aHUS TMA (CFI mutation) | 1 |
| interstitial nephritis | 1 |
| rare (nephropathic cystinosis, lymphangioleiomyomatosis) | 2 |
| unclear | 1 |
| sum | 19 |

## Notes

### Competing Interest Statement

The authors have declared no competing interest.

